# A pyro-phosphodegron controls MYC polyubiquitination to regulate cell survival

**DOI:** 10.1101/2020.02.12.945634

**Authors:** Padmavathi Lolla, Akruti Shah, C.P. Unnikannan, Vineesha Oddi, Rashna Bhandari

## Abstract

The transcription factor MYC regulates cell survival and growth, and its level is tightly controlled in normal cells. Here, we report that serine pyrophosphorylation – an enigmatic posttranslational modification triggered by inositol pyrophosphate signaling molecules – controls MYC levels via regulated protein degradation. We find that endogenous MYC is stabilized and less polyubiquitinated in cells with reduced inositol pyrophosphates. We show that the inositol pyrophosphate 5-IP_7_ transfers its high-energy beta phosphate moiety to pre-phosphorylated serine residues in the central PEST domain of MYC. Pyrophosphorylation of MYC promotes its interaction with the E3 ubiquitin ligase FBW7, thereby enhancing MYC polyubiquitination and degradation. FBW7 can bind directly to the PEST domain of MYC in a pyrophosphorylation-dependent manner. A stabilized, pyrophosphorylation-deficient form of MYC increases cell death during growth stress in untransformed cells, and promotes cell proliferation in response to mitogens. Thus, control of MYC stability through a novel pyro-phosphodegron provides unexpected insight into the regulation of cell survival in response to environmental cues.

## Introduction

Inositol pyrophosphates are derivatives of *myo*-inositol characterized by the presence of one or more diphosphates in addition to monophosphates around the inositol ring (1). These small molecules are found in all eukaryotic cells, and are involved in the regulation of many cellular processes, including energy and phosphate metabolism, DNA repair, ribosome biogenesis, vesicle trafficking and insulin secretion (1–5). In mammals, the most abundant inositol pyrophosphate is 5-diphosphoinositol pentakisphosphate (5-PP-IP5 or 5-IP_7_), which is synthesized from inositol hexakisphosphate (IP_6_) by IP6 kinases. There are three isoforms of IP6 kinase in mammals – IP6K1, 2, and 3 (6–8). Inositol pyrophosphates are capable of transferring their high-energy β-phosphate moiety to pre-phosphorylated serine residues (9–11), thus pyrophosphorylating proteins to influence their function *in vivo* (12–15). Acidophilic protein kinases such as casein kinase 2 (CK2), preferentially phosphorylate Ser residues in the vicinity of Glu/Asp residues, priming them for pyrophosphorylation (10). Ser residues occurring in mobile disordered regions that are rich in Asp/Glu residues may be good substrates for pyrophosphorylation (11, 14, 15).

MYC (c-MYC) is a transcription factor that accelerates cell growth and proliferation by regulation of genes involved in glucose and amino acid metabolism, ribosomal and mitochondrial biogenesis, protein synthesis, and cell cycle progression (16) among others. MYC levels are upregulated in a variety of cancer types (17), but in normal cells, the level of this oncoprotein is tightly regulated (18). MYC protein is short-lived and has a central PEST domain (enriched in Pro, Glu/Asp, Ser, Thr residues) (19) which contributes to its instability (20). This disordered region (21) has many Ser residues that could be potential targets for pyrophosphorylation by inositol pyrophosphates.

Here we show that 5-IP_7_ mediated pyrophosphorylation of Ser residues lying within the PEST domain of MYC regulates its turnover. We observed a two-fold increase in MYC half-life in cells depleted for 5-IP_7_. Upon pyrophosphorylation, the PEST domain of MYC can bind to the E3 ubiquitin ligase FBW7, resulting in increased polyubiquitination and degradation of MYC. Stabilization of MYC upon a reduction in its pyrophosphorylation correlates with reduced cell survival under stress, and increased cell proliferation upon growth stimulation.

## Results

### Loss of IP6K1 leads to increased MYC stability

It has been shown that mouse embryonic fibroblasts (MEFs) derived from IP6K1 knockout mice display a 70% decrease in the level of 5-IP_7_ (22). To determine if 5-IP_7_ regulates MYC, we compared the levels of endogenous mouse MYC (mMYC) protein in MEFs derived from *Ip6k1*^+/+^ and *Ip6k1*^−/−^ mice. Steady state mMYC protein levels were approximately two fold higher in *Ip6k1*^−/−^ MEFs compared to their respective *Ip6k1*^+/+^ counterparts (Fig. 1A). MYC has a reported half-life of 20-30 min (20), and in keeping with this, we observed that the half-life of mMYC in *Ip6k1*^+/+^ MEFs is 25 min (Fig. 1B and C). mMYC was approximately twice as stable in *Ip6k1*^−/−^ MEFs compared to *Ip6k1*^+/+^ MEFs, suggesting that IP6K1 downregulates the stability of MYC under normal growth conditions. MYC is degraded mainly by the ubiquitin-proteasome system (18). We found that mMYC is less poly-ubiquitinated in *Ip6k1*^−/−^ MEFs compared with *Ip6k1*^+/+^ MEFs (Fig. 1D), suggesting that the increased half-life of mMYC in cells lacking IP6K1 is due to a decrease in ubiquitin-dependent proteasomal degradation.

**Figure 1.**
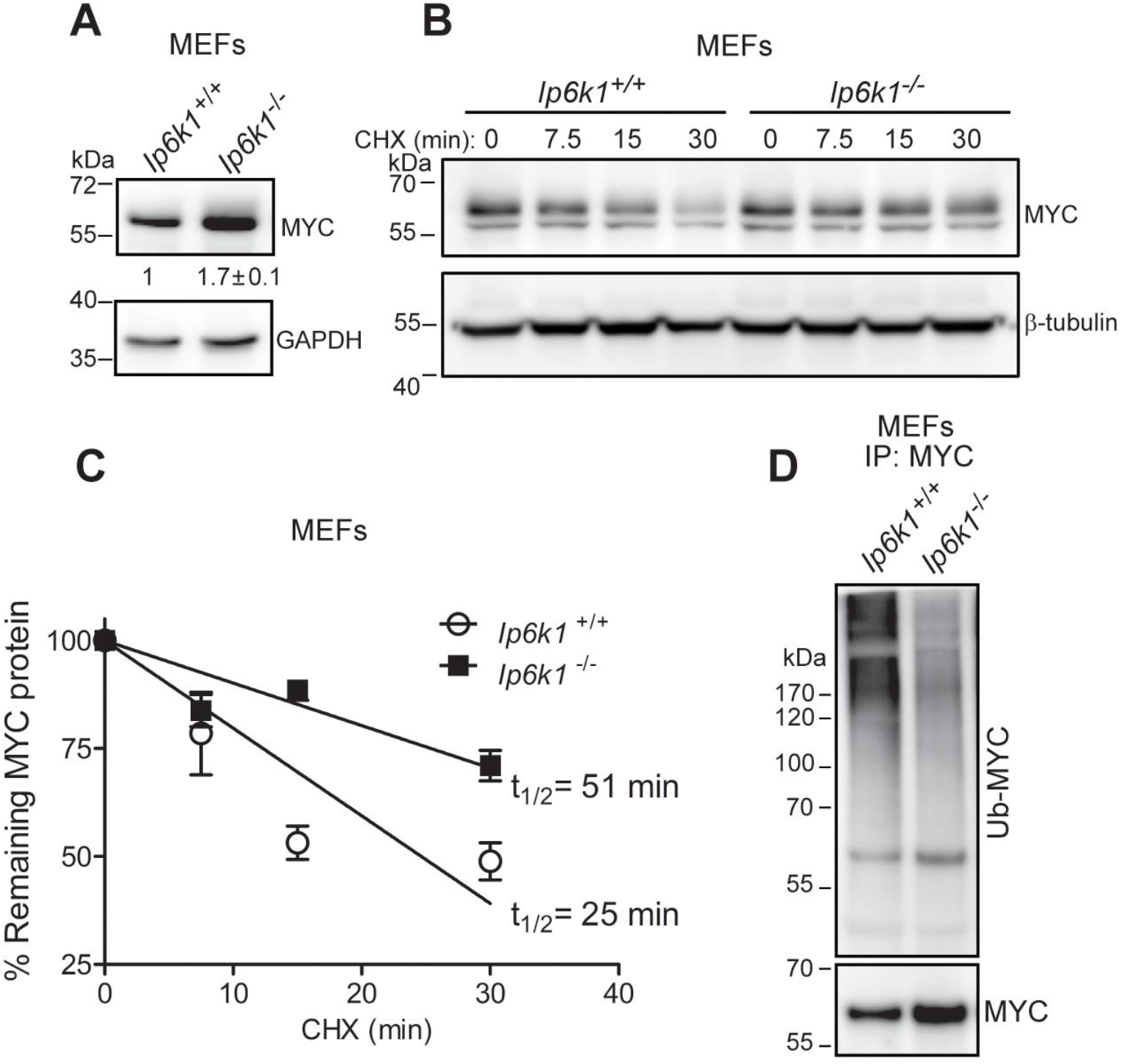
MYC protein is stabilized in *Ip6k1*^−/−^ MEFs. **(A)** Immunoblot showing steady state MYC protein levels in MEFs from *Ip6k1*^+/+^ and *Ip6k1*^−/−^ mice. Numbers show mean fold change±SEM (N=3) in the level of MYC in *Ip6k1*^−/−^ over *Ip6k1*^+/+^ MEFs **(B)** Representative immunoblot for half-life measurement of MYC in *Ip6k1*^+/+^ and *Ip6k1*^−/−^ MEFs (N=3). Cells were treated with 100 µg/mL cycloheximide (CHX) for the indicated time, lysed and immunoblotted. **(C)** MYC levels (normalized to loading control) at different time points after treatment with CHX, as in (B), were plotted as a percentage of MYC in untreated cells, and half-life was calculated by linear regression analysis. Data are mean ± SEM (N=3). **(D)** Representative immunoblot showing polyubiquitination of endogenous MYC (Ub-MYC) from *Ip6k1*^+/+^ and *Ip6k1*^−/−^ MEFs. Cells were treated with 20µM MG132 for 2 h, lysed, immunoprecipitated with anti-MYC antibody, and immunoblotted with an anti-ubiquitin antibody or anti-MYC antibody (N=3).

### MYC undergoes pyrophosphorylation by 5-IP_7_

We wondered whether the IP6K1-dependent decrease in MYC stability is due to pyrophosphorylation of MYC by 5-IP_7_. To test for MYC pyrophosphorylation, we transiently expressed C-terminally V5 epitope-tagged mMYC in HEK293T cells and incubated the immunoprecipitated protein with 5[β-^32^P]IP_7_ (Fig. 2A). We found that this exogenously expressed mMYC protein does indeed get pyrophosphorylated by 5-IP_7_ *in vitro*. To determine whether endogenous mMYC is pyrophosphorylated *in vivo*, we conducted a ‘back-pyrophosphorylation’ assay (9, 12, 15, 23). In this assay, a protein that is already pyrophosphorylated *in vivo* shows reduced acceptance of radiolabeled phosphate from 5-IP_7_ *in vitro*, whereas the same target protein isolated from cells with reduced 5-IP_7_ has diminished *in vivo* pyrophosphorylation, and shows augmented pyrophosphorylation *in vitro* (see schematic in Fig. 2B). mMYC protein isolated from *Ip6k1*^−/−^ MEFs showed robust incorporation of radiolabeled phosphate from 5-IP_7_, whereas mMYC isolated from *Ip6k1*^+/+^ MEFs showed a marginal level of radiolabeling, reflecting the high levels of *in vivo* pyrophosphorylation of mMYC in *Ip6k1*^+/+^ MEFs (Fig. 2B). Human MYC (hMYC) shows 91% sequence identity with mMYC. We confirmed that hMYC is also pyrophosphorylated by 5-IP_7_. hMYC, expressed as a fusion to GST in bacteria, was first phosphorylated by CK2, the Ser/Thr kinase that pre-phosphorylates Ser residues which can undergo subsequent pyrophosphorylation (10, 24) (Fig. S1A). GST-hMYC that was pre-phosphorylated by CK2 using unlabeled ATP, subsequently underwent pyrophosphorylation upon incubation with 5[β-^32^P]IP_7_ (Fig. S1B).

**Figure 2.**
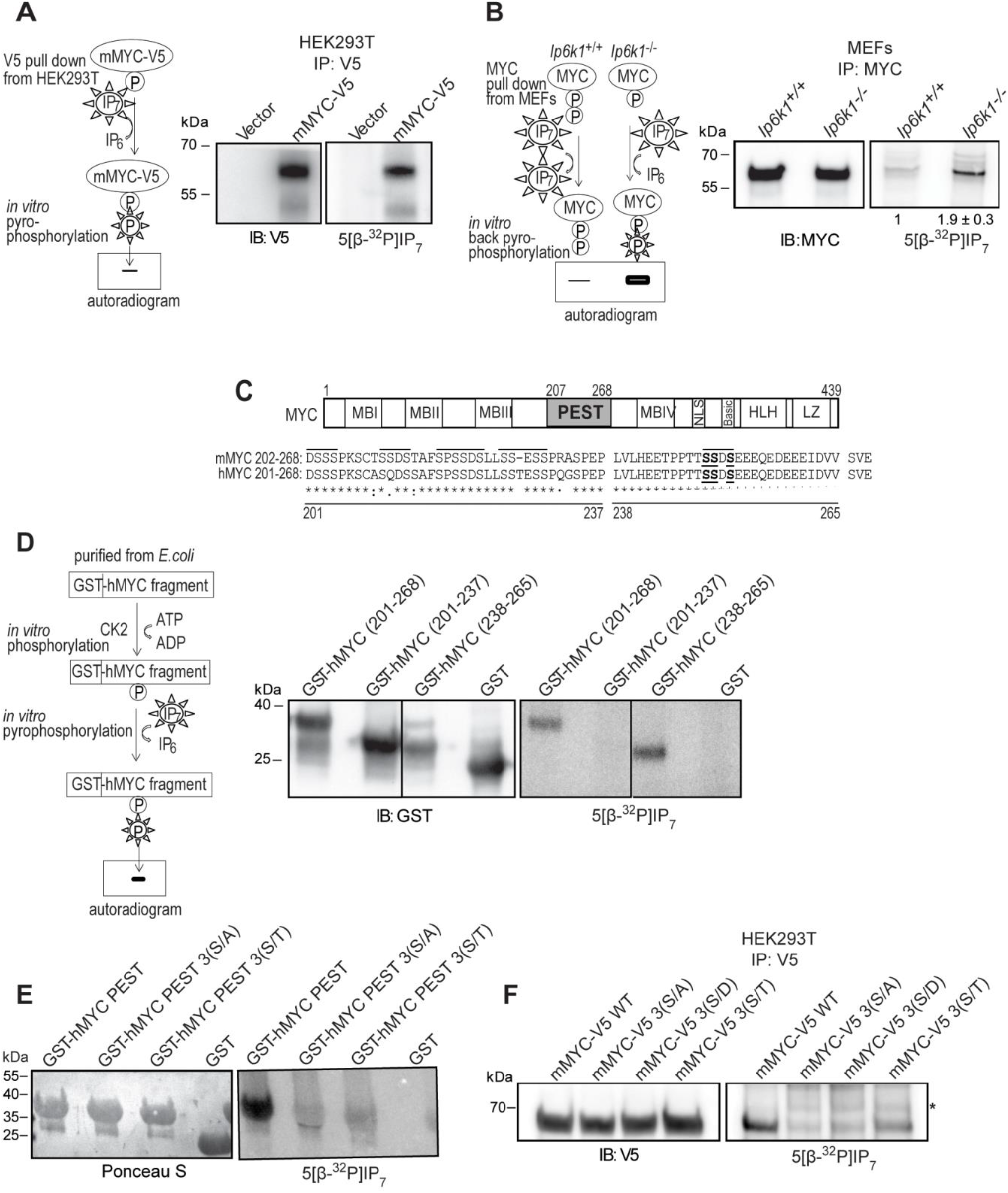
MYC is pyrophosphorylated by 5-IP_7_ in its central PEST domain. **(A)** Mouse MYC tagged C-terminally with V5 (mMYC-V5) was transiently expressed in HEK293T cells, immunoprecipitated, incubated with 5[β-^32^P]IP_7_, resolved by NuPAGE, and transferred to a PVDF membrane (shown schematically on the left). Representative images show autoradiography to determine pyrophosphorylation (right) and immunoblotting with a V5-tag antibody (left) (N=3). Cells transfected with pCDNA3.1(+) plasmid (vector) served as a negative control. **(B)** Schematic on the left describes the back-pyrophosphorylation assay (23). Endogenous MYC from *Ip6k1*^+/+^ and *Ip6k1*^−/−^ MEFs was immunoprecipitated and pyrophosphorylated in presence of 5[β-^32^P]IP_7_ as in **A**. Representative images show autoradiography to detect pyrophosphorylation (right) and immunoblotting to detect MYC (left). Numbers show mean fold change±SEM in the extent of back-pyrophosphorylation in *Ip6k1*^−/−^ over *Ip6k1*^+/+^ MEFs (N=3). **(C)** Domain map of MYC. Mouse MYC (mMYC) and human MYC (hMYC) show 91% identity in their PEST domains (protein-protein BLAST); all acidic Ser motifs (overlined) are conserved; pyrophosphorylated Ser cluster is in bold and underlined. **(D)** Purified, GST-tagged hMYC (GST-hMYC) PEST domain fragments were phosphorylated by CK2 in presence of unlabeled ATP, and then pyrophosphorylated by 5[β-^32^P]IP_7_ (shown schematically on the left). Representative images show autoradiography to detect pyrophosphorylation (right) and immunoblotting with a GST antibody (left) (N=2). Line indicates removal of non-essential lanes from a single original gel. Start and end amino acid numbers for hMYC fragments are indicated in brackets. **(E)** Purified, GST-hMYC PEST domain (aa 201-268) corresponding to the native sequence, or with three Ser residues (249/250/252) mutated to Ala (3(S/A)) or Thr (3(S/T)), were treated as described in **D** (N=2). Proteins were detected by Ponceau S staining of the membrane. (**F**) Full-length mMYC-V5 protein (WT) or its mutant forms, Ser 249/250/252 replaced with Ala (3(S/A)), Asp (3(S/D)) or Thr (3(S/T)), were transiently expressed in HEK293T cells and treated as in **A** (N=2). Native mMYC shows pyrophosphorylation as in **A**, mMYC 3(S/T) shows faint pyrophosphorylation, whereas mMYC 3(S/A) and 3(S/D) are of the same intensity as non-specific bands (marked by an asterisk).

### MYC pyrophosphorylation is localized to Ser residues within its PEST domain

We hypothesized that the most likely site of pyrophosphorylation on MYC is its PEST domain, as this region possesses several Ser residues in the vicinity of Glu/Asp residues, and its deletion has been shown to stabilize MYC (20). The hMYC PEST domain spanning amino acid residues (aa) 207-268 (20), aligns with aa208-268 in mMYC (Fig. 2C). There is a short acidic serine motif immediately N-terminal to the PEST sequence, which we included in our search for the site of pyrophosphorylation on MYC. An alignment of aa202-268 of mMYC with aa201-268 of hMYC revealed conservation of all the acidic serine clusters (Fig. 2C). GST-hMYC aa201-268 was phosphorylated by CK2 in the presence of [γ^32^-P]ATP (Fig. S1C). An earlier report using synthetic peptide fragments had shown that CK2 can phosphorylate hMYC in the region between residues 240-262 (25). To narrow down the CK2 target sites in the PEST domain, we divided this region into two fragments – hMYC aa201-237 and aa238-265 (Fig. 2C), and examined these GST fusion peptides for phosphorylation by CK2. Of these, only the protein fragment aa238-265, containing the previously reported CK2 target site (25), was phosphorylated (Fig. S1C). To examine 5-IP_7_ dependent pyrophosphorylation of the MYC PEST domain, we subjected the same protein fragments to CK2 phosphorylation using unlabeled ATP, and subsequently incubated them with 5[β-^32^P]IP_7_ (24). The GST fused hMYC PEST domain aa201-268 underwent pyrophosphorylation under these conditions (Fig. 2D). The PEST fragment aa238-265, which was phosphorylated by CK2, also showed pyrophosphorylation by 5-IP_7_, whereas aa201-237 was not pyrophosphorylated (Fig. 2D), thus narrowing down the 5-IP_7_ target site to the C-terminal region of the MYC PEST domain. This region contains a cluster of three Ser residues, S249, S250 and S252 (Fig. 2C, bold and underlined), in the vicinity of Glu and Asp residues. Previous studies have shown that multiple Ser residues within an acidic serine cluster can undergo pyrophosphorylation, and that pyrophosphorylation can be eliminated by replacing all Ser residues in the cluster with Ala (10, 13, 14). We therefore generated a mutant version of the hMYC PEST domain called 3(S/A), in which Ser 249, 250 and 252 were all replaced with Ala. The loss of these three Ser residues nearly eliminated CK2 phosphorylation of the MYC PEST domain (Fig. S1D), suggesting that the major site of CK2 phosphorylation lies within this Ser cluster. Replacing these Ser residues with Ala also led to a substantial reduction in 5[β-^32^P]IP_7_-mediated pyrophosphorylation (Fig. 2E), thus localizing the site of pyrophosphorylation on MYC to one or more of Ser residues 249/250/252. We also examined the effect of replacing these three Ser residues with Thr on CK2-mediated phosphorylation and on 5-IP_7_ mediated pyrophosphorylation of hMYC PEST domain. CK2 was able to phosphorylate the 3(S/T) mutant PEST sequence to the same extent as the native PEST sequence (Fig. S1D). However, despite its robust pre-phosphorylation by CK2, the 3(S/T) mutant PEST domain showed weak pyrophosphorylation (Fig. 2E). This data suggests that phosphoThr does not accept the β-phosphate from 5-IP_7_ with the same efficiency as phosphoSer, despite being in the same sequence context.

### Pyrophosphorylation of MYC promotes its polyubiquitination

To determine the effect of phosphorylation or pyrophosphorylation of the PEST domain on the stability of MYC, we generated three mutant forms of full length mMYC – 3(S/A), 3(S/T), and 3(S/D), in which Ser 249, 250 and 252 were replaced with (i) Ala, to prevent phosphorylation and pyrophosphorylation, (ii) Thr, to allow normal phosphorylation/dephosphorylation but reduced pyrophosphorylation, or (iii) Asp, to mimic constitutive phosphorylation but no pyrophosphorylation. Replacing Ser 249/250/252 with Ala or Asp in full length mMYC led to a near complete elimination of 5[β-^32^P]IP_7_-mediated pyrophosphorylation, confirming that this is indeed the major pyrophosphorylation region on MYC (Fig. 2F). As in the case of hMYC PEST domain (Fig. 2E), substituting Thr in place of Ser in full length mMYC resulted in a substantially reduced level of pyrophosphorylation (Fig. 2F). Transient expression of low amounts of these three forms of mMYC in HEK293T cells revealed that the level of all three mutant proteins was approximately 70% higher compared with the wild type (WT) protein (Fig. 3A). To examine the stability of these mMYC mutants, we monitored their levels for up to 3 h after treating cells with cycloheximide. All three pyrophosphorylation-deficient mutant forms of mMYC showed an approximately two-fold greater stability than WT mMYC (Fig. 3B and C), correlating with the two-fold increase in half-life of endogenous mMYC observed in *Ip6k1*^−/−^ MEFs (Fig. 1B and C). An earlier study has reported that CK2 protects MYC from proteasomal degradation, but the site(s) of CK2 mediated phosphorylation on MYC responsible for this effect were not identified (26). Our data revealed no significant difference in half-life between the non-phosphorylated 3(S/A) mutant and the phosphorylated 3(S/T) or phosphomimic 3(S/D) mutant forms, suggesting that Ser 249/250/252 may not be the CK2 phosphosites responsible for increasing MYC stability. We then examined whether the loss of PEST domain pyrophosphorylation alters the extent of MYC poly-ubiquitination. The pyrophosphorylation-deficient mutant forms of mMYC showed reduced poly-ubiquitination compared to WT mMYC (Fig. 3D), correlating with the difference in their half-lives (Fig. 3B and C). Overall, our results suggest that phosphorylation of Ser 249/250/252 does not affect MYC stability, and that *pyro*phosphorylation at this site is necessary to maintain MYC turnover in cells.

**Figure. 3.**
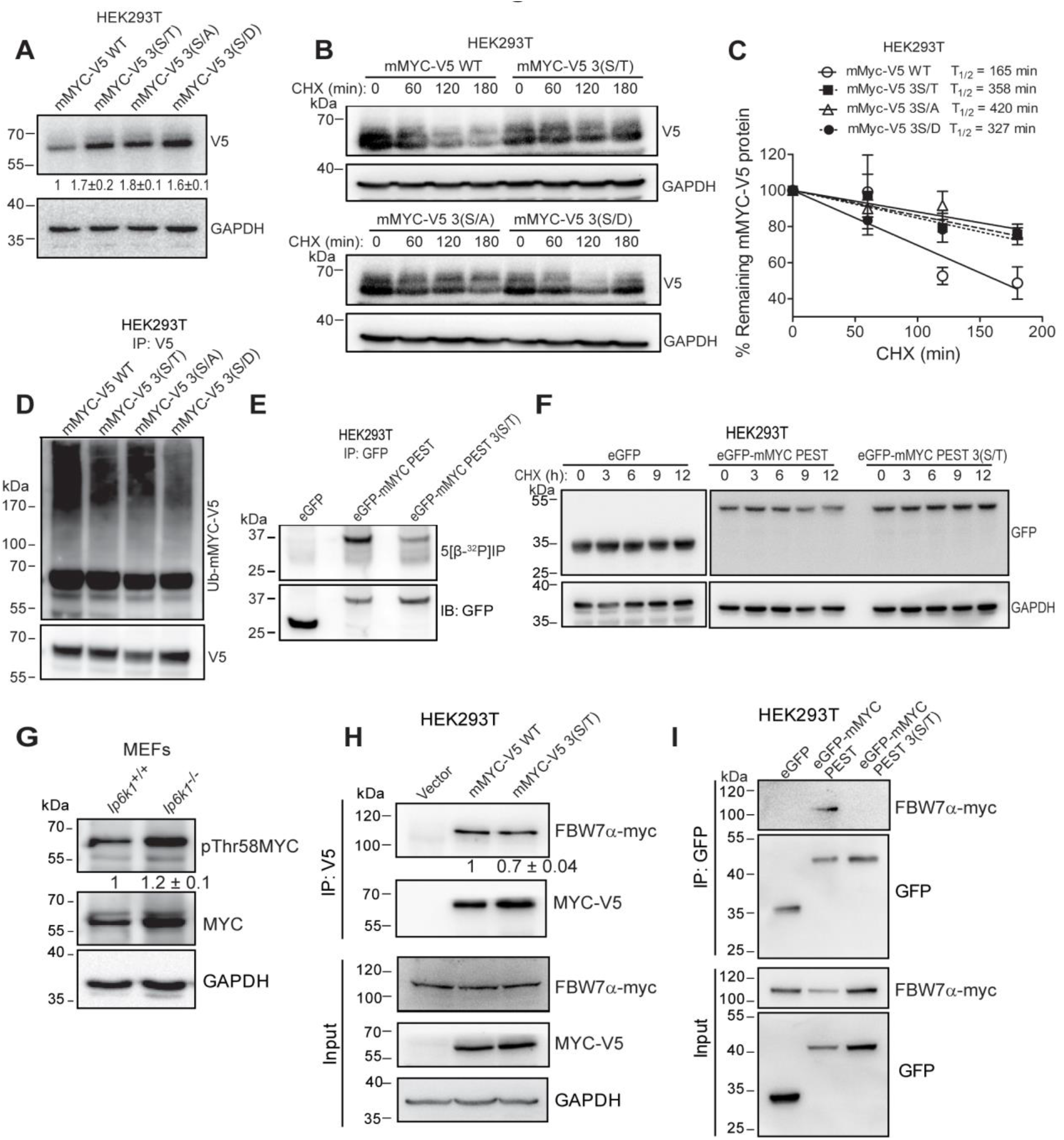
PEST domain pyrophosphorylation promotes F-box protein FBW7α binding and polyubiquitination of MYC. **(A)** Full-length mMYC-V5 protein and its mutant forms were transiently expressed in HEK293T cells. Numbers show mean fold change±SEM in expression level of individual mutants over the native protein (N=3). (**B, C**) Representative immunoblot for half-life measurement of mMYC-V5 forms. Cells were treated with 250 µg/mL CHX for the indicated time, and half-life was determined as described in **Fig. 1C** (N=3). (**D**) mMYC-V5 WT and its mutant forms were transiently expressed in HEK293T cells and treated as in Fig 1D, immunoprecipitated with anti-V5 antibody and immunoblotted with an anti-ubiquitin antibody (Ub-mMYC-V5) or anti-V5 antibody (N=3). (**E**) eGFP fused C-terminally with native or 3(S/T) mutant form of mMYC PEST domain (aa 202-268) was transiently expressed in HEK293T cells, immunoprecipitated, incubated with 5[β-^32^P]IP_7_, resolved by NuPAGE, and transferred to a PVDF membrane, as described in **Fig. 2A**. Representative images show autoradiography to determine pyrophosphorylation (top) and immunoblotting with a GFP antibody (bottom) (N=2). (**F**) Representative immunoblots to assess the stability of eGFP, eGFP-mMYC PEST, or eGFP-mMYC PEST 3(S/T). HEK293T cells transiently expressing the different forms of eGFP were treated with 100 µg/mL cycloheximide (CHX) for the indicated time, lysed and immunoblotted to detect eGFP. GAPDH levels were monitored as a loading control (N=3). **(G)** Representative immunoblot showing the extent of phosphorylation at Thr58 and expression levels of endogenous MYC in *Ip6k1*^+/+^ and *Ip6k1*^−/−^ MEFs. Numbers show mean fold change±SEM in Thr58 phosphorylation normalized to MYC levels in *Ip6k1*^−/−^ over *Ip6k1*^+/+^ (N=3). **(H)** Representative immunoblots of interaction between mMYC and FBW7α. mMYC-V5 WT and pyrophosphorylation-deficient mutant mMYC-V5 3(S/T) were transiently co-expressed with c-Myc epitope-tagged FBW7α in HEK293T cells, immunoprecipitated with anti-V5 antibody, and probed to detect c-Myc epitope. The level of coimmunoprecipitated FBW7α was normalized to the level of immunoprecipitated MYC. Numbers show mean fold change±SEM in the extent of coimmunoprecipitation of FBW7α with 3(S/T) MYC compared with WT MYC (N=3) (**I**) Representative immunoblots of FBW7α interaction with eGFP fused with mMYC PEST domain. eGFP-mMYC PEST or eGFP-mMYC PEST 3(S/T) were transiently co-expressed with c-Myc epitope tagged FBW7α in HEK293T cells, immunoprecipitated with anti-GFP antibody, and probed to detect c-Myc epitope. No coimmunoprecipitation of FBW7α was detected with eGFP fused with pyrophosphorylation-deficient mMYC PEST 3(S/T) (N=3).

### A ‘pyro-phosphodegron’ is essential for binding of the MYC PEST domain to FBW7 E3 ligase

Next, we probed the molecular mechanism by which pyrophosphorylation in the PEST domain leads to MYC destabilization. It has been shown that the stability of GFP can be significantly lowered by C-terminally fusing it to the PEST domain of mouse ornithine decarboxylase (27, 28). We fused the WT and 3(S/T) versions of mMYC PEST domain (aa202-268) to the C-terminus of enhanced GFP (eGFP). Incubation of eGFP-tagged WT mMYC PEST domain with radiolabeled 5-IP_7_ led to its pyrophosphorylation, whereas eGFP fused with the 3(S/T) mutant mMYC PEST domain showed significantly reduced pyrophosphorylation (Fig. 3E). eGFP alone was not pyrophosphorylated. Treatment of cells with cycloheximide for up to 12 h did not lead to any change in the level of native eGFP, which is known to be a highly stable protein (28). However, there was an appreciable decrease in eGFP fused to WT mMYC PEST domain over the same period of time (Fig. 3F). Fusion with the 3(S/T) mutant version of mMYC PEST domain did not significantly alter the stability of eGFP (Fig. 3F), indicating that pyrophosphorylation of the MYC PEST domain is important for its effect on protein stability.

Several E3 ligases have been shown to contribute to the poly-ubiquitination and proteasome-mediated degradation of MYC (18). We however focused on FBW7, a component of the Skp-Cullin-F box (SCF) ubiquitin ligase complex, which is known to bind phosphorylated Thr/Ser and Pro containing sequences referred to as a phosphodegron (29, 30). FBW7 binding to MYC has been shown to require GSK3β-mediated MYC phosphorylation at Thr58 (20, 31). The kinase activity of GSK3β can be stimulated by binding to IP6K1 *in vitro*, and this effect is independent of IP6K1 catalytic activity (32). Therefore, loss of IP6K1 could lower GSK3β activity, reducing Thr58 phosphorylation, and thereby increasing MYC stability. To rule out any 5-IP_7_-independent effect on MYC stability via regulation of GSK3β, we examined the level of MYC Thr58 phosphorylation in MEFs lacking IP6K1. When normalized to MYC levels, there was no significant difference in the extent of Thr58 phosphorylation between *Ip6k1*^+/+^ and *Ip6k1*^−/−^ MEFs, indicating that the effect of IP6K1 on MYC stability is not via GSK3β (Fig. 3G). To check whether FBW7 binding to MYC is influenced by PEST domain pyrophosphorylation, we compared the extent of FBW7α binding to WT and 3(S/T) mMYC. We observed a 30% decrease in FBW7α binding to the pyrophosphorylation-deficient 3(S/T) mutant form of mMYC compared with the WT form (Fig. 3H). This interaction of FBW7α with full length mMYC reflects FBW7α binding at multiple possible sites on MYC, including its previously demonstrated binding site at the Thr58 phosphodegron (31). To specifically examine the binding between FBW7α and the MYC PEST domain, we co-expressed FBW7α with eGFP-tagged WT or 3(S/T) versions of the mMYC PEST domain. Although FBW7α did not bind to native eGFP, it showed a robust interaction with eGFP fused to the WT mMYC PEST domain (Fig. 3I). However, eGFP fused to the 3(S/T) mutant mMYC PEST domain showed no interaction with FBW7α. This indicates that compromising pyrophosphorylation abolishes the ability of the mMYC PEST domain to bind FBW7α. These data suggest that the SSDS motif (aa249-252) in the PEST domain of MYC (Fig. 2C) constitutes a novel “pyro-phosphodegron”, which undergoes phosphorylation by acidophilic protein kinases such as CK2, and subsequent pyrophosphorylation by inositol pyrophosphates to permit binding by FBW7, polyubiquitination and degradation of MYC.

### Pyrophosphorylation fine tunes MYC levels to regulate cell fate in response to environmental cues

Finally, we wanted to examine the biological consequences of pyrophosphorylation-dependent changes in MYC stability. We stably expressed either WT or 3(S/T) mMYC in the HO15.19 *myc*^−/−^ Rat1 immortalized fibroblast cell line, in which both copies of endogenous *Myc* gene have been disrupted (33). In line with its increased stability, 3(S/T) mMYC expressed in *Myc*-null rat fibroblasts showed a two-fold increase in steady state levels compared to WT mMYC (Fig. 4A). We examined anchorage independent growth and clonogenic capacity in Rat1 cells carrying the two forms of MYC. Compared with *Myc*-null cells, expression of mMYC permitted Rat1 fibroblasts to form single-cell derived colonies on soft-agar (Fig. 4B and C). Interestingly, cells expressing the more stable 3(S/T) mutant form of mMYC showed a marked reduction in the number of colonies compared to cells carrying WT mMYC. Similarly, the clonogenic capacity of 3(S/T) mMYC expressing cells was lower than that of cells with WT mMYC (Fig. 4D and E). It has been shown that MYC harbors latent pro-apoptotic activity which is manifested when MYC is elevated, and is especially apparent when cells are subjected to limitation of nutrients and growth factors (34, 35). In Rat1 immortalized fibroblasts, increased MYC expression has been correlated with higher cell death upon serum depletion (36). We examined the level of cell death in HO15.19 *myc*^−/−^ Rat1 lines carrying WT or 3(S/T) mMYC, grown in normal *vs* reduced serum conditions. Cells expressing 3(S/T) mMYC showed elevated cell death when grown in serum depleted medium, whereas the level of cell death in cells expressing WT mMYC was unaffected by serum depletion (Fig. 4F).

**Fig. 4.**
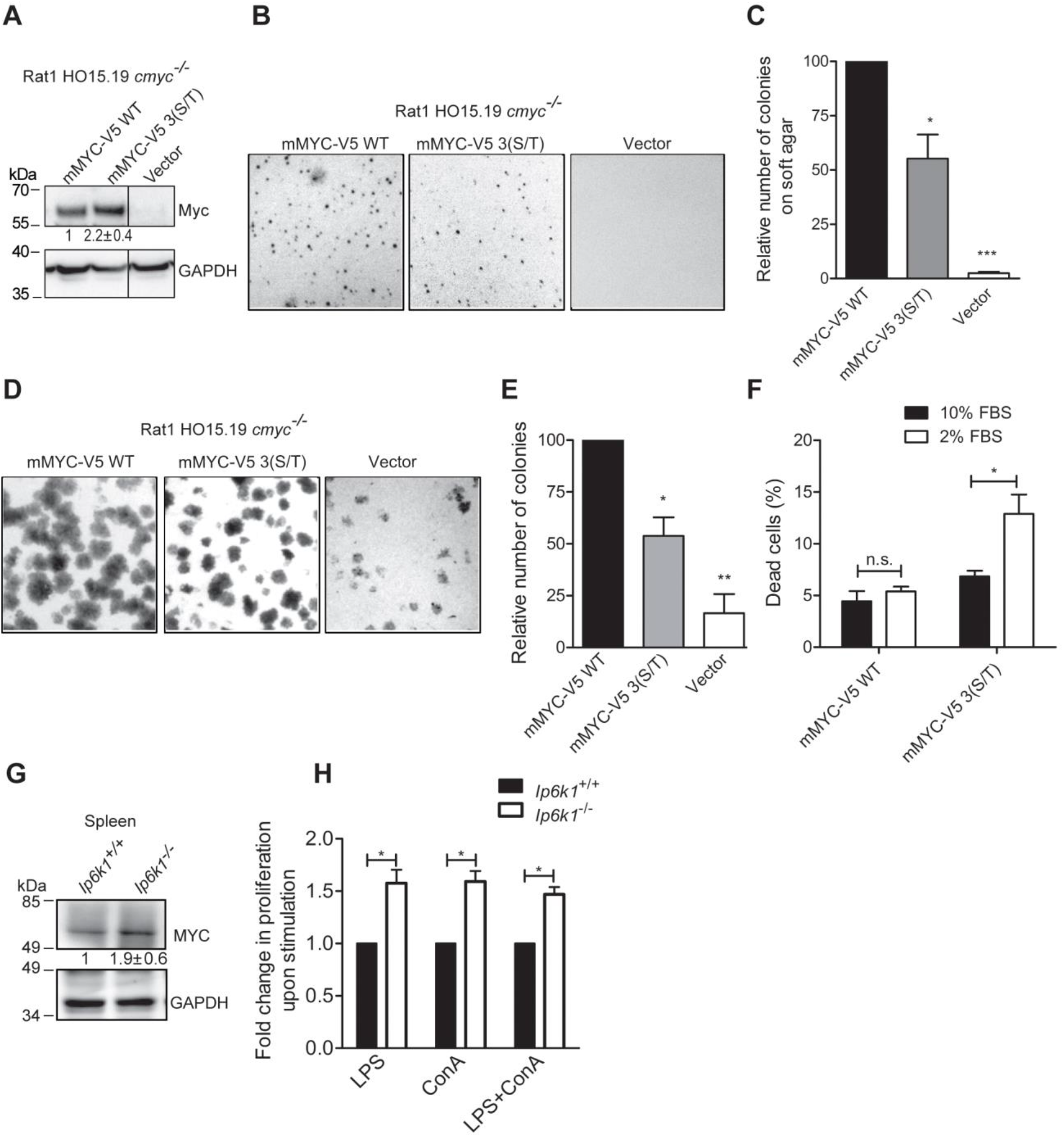
MYC stabilization upon loss of pyrophosphorylation influences cell survival and proliferation. (**A**) Immunoblot showing steady state levels of mMYC-V5 WT and its pyrophosphorylation-deficient mutant form mMYC-V5 3(S/T), stably expressed in Rat1 HO15.19 *cmyc*^−/−^ cells. Numbers show mean fold change±SEM in level of the mutant form over the WT protein (N=3). (**B)** Representative images showing anchorage-independent growth of Rat1 HO15.19 *cmyc*^−/−^ cells expressing mMYC-V5 WT or 3(S/T). Cells transfected with pBABE-Puro plasmid (vector) served as a control. (**C**) Images from **B** were quantified using Fiji software and colony numbers were plotted relative to mMYC-V5 WT (data are mean±SEM, N=3). (**D**) Representative images showing colony formation by Rat1 HO15.19 *cmyc*^−/−^ cells expressing mMYC-V5 WT, 3(S/T), or vector alone. (**E**) Images from **D** were quantified using Fiji software and colony numbers were plotted relative to mMYC-V5 WT (data are mean±SEM, N=3). **(F)** Rat1 HO15.19 *cmyc*^−/−^ cells expressing mMYC-V5 WT or 3(S/T) were cultured in either serum rich (10% FBS) or depleted (2% FBS) medium for 48 h, dead cells were stained with propidium iodide, and percentage of dead cells was estimated by flow cytometry (N=3). **(G)** Immunoblot showing steady state MYC protein levels in spleens from *Ip6k1*^+/+^ and *Ip6k1*^−/−^ mice. Numbers show mean fold change±SEM (N=3) in the level of MYC in *Ip6k1*^−/−^ over *Ip6k1*^+/+^. **(H)** Mitogenic response of lymphocytes enriched from spleens of either *Ip6k1*^+/+^ or *Ip6k1*^−/−^ mice measured by the MTT colorimetric assay. Absorbance value for each mitogen treatment condition was normalized to the respective mock treated sample. Proliferation of *Ip6k1*^−/−^ lymphocytes was expressed as a fold change over *Ip6k1*^+/+^ lymphocytes for each mitogen treatment condition (data are mean±SEM, N=3). *P* values are from a one-sample *t*-test (**C**, **E and H**), or a two-tailed unpaired Student’s *t*-test (**F**) (\**P*≤0.05; \*\**P*≤0.01; \*\*\**P*≤0.001; n.s. not significant, P >0.05).

To investigate the effect of MYC pyrophosphorylation on cell proliferation, we turned to spleen tissue, as spleen resident T and B lymphocytes are known to express high levels of MYC during proliferation (37). Steady state MYC levels in spleen tissue from *Ip6k1*^−/−^ mice showed a two-fold increase when compared to *Ip6k1*^+/+^ mice (Fig. 4G). We enriched lymphocytes from spleens of *Ip6k1*^+/+^ and *Ip6k1*^−/−^ mice, and subjected them to stimulation by the mitogens lipopolysaccharide (LPS) and concanavalin A (ConA), which are known to stimulate MYC-driven proliferation of B and T lymphocytes respectively (37). Lymphocytes derived from *Ip6k1*^−/−^ mice displayed an approximately 50% increase in proliferation when compared to their *Ip6k1*^+/+^ counterparts, upon stimulation with either LPS, ConA, or LPS and ConA (Fig. 4H). Together these data reveal that cellular outcomes of pyrophosphorylation-deficient MYC stabilization are context dependent – leading to an increase in cell death under conditions of cellular stress, but accentuating the proliferative capacity of cells in an environment conducive to growth.

## Discussion

Our work uncovers 5-IP_7_ as a new player in the biology of MYC. We have shown that 5-IP_7_-mediated serine pyrophosphorylation within the PEST domain of this oncoprotein facilitates its polyubiquitination and turnover. In this way, alteration in cellular levels of inositol pyrophosphates could fine tune MYC in response to environmental cues. Earlier work on inositol pyrophosphate dependent serine pyrophosphorylation has shown that this post-translational modification regulates housekeeping cellular functions including glycolysis, ribosome synthesis, and cargo binding to motor proteins (12–15). Our current study reveals a role for serine pyrophosphorylation in the regulation of cell survival and proliferation.

We have coined the term “pyro-phosphodegron” to describe the sequence SSDS (aa249-252) within the PEST domain of MYC, which binds to FBW7 upon serine pyrophosphorylation by 5-IP_7_. The sequence TPPTT (aa244-248; Fig. 2C), lying immediately N-terminal to the MYC pyro-phosphodegron, has similarity to the N-terminal MYC phosphodegron (aa58-62) and has been predicted to bind FBW7 (38). Thr244 is endogenously phosphorylated, and replacing it with Ala results in stabilization of MYC (38). Together with our data, this suggests that the sequence TPPTTSSDS (aa244-252) forms a contiguous phospho/pyrophospho-degron that controls MYC stability via binding to FBW7. FBW7 is known to function as a dimer and each protomer in dimeric FBW7 can interact with identical sites on a homodimeric substrate, or different sites on a monomeric substrate protein (39, 40). Such cooperative binding has been shown to increase binding affinity and promote greater turnover of the target protein (39–41). Loss of FBW7 dimerization has been reported to lower the turnover of endogenous MYC (40), suggesting that there may be two binding sites on MYC for dimeric FBW7. However, until now, MYC has only been shown to bind FBW7 via its N-terminal Thr58 phosphodegron. Our demonstration of a second binding site for FBW7 in the MYC PEST domain suggests that MYC is capable of binding cooperatively to FBW7 (Fig. S2). This mode of binding between MYC and dimeric FBW7 would allow for greater binding affinity, enhanced polyubiquitination, and flexibility in signaling pathways that can promote MYC turnover.

It is likely that PEST domain pyrophosphorylation on other short-lived proteins regulates their interaction with FBW7 and perhaps other E3 ligases, facilitating their turn over. In this way, serine pyrophosphorylation by inositol pyrophosphates may emerge as a key posttranslational modification in the regulation of protein stability, thereby influencing several cellular functions.

## Materials and Methods

Details of experimental procedures and statistical analysis methods used in this study are described in Supplementary Information.

### Mice

All animal experiments were approved by the Institutional Animal Ethics Committee (Protocol number PCD/CDFD/02 – version 2), and were performed in compliance with guidelines provided by the Committee for the Purpose of Control and Supervision of Experiments on Animals, Ministry of Environment, Forest, and Climate Change, Government of India. Mice *(Mus musculus*, strain C57BL/6) used for this study were housed in the Centre for DNA Fingerprinting and Diagnostics animal facility located within the premises of Vimta Labs, Hyderabad. The *Ip6k1* gene knockout mouse was generated as previously described (22). *Ip6k1*^+/+^ and *Ip6k1*^−/−^ littermates were produced by breeding *Ip6k1*^+/−^ mice.

### Phosphorylation and pyrophosphorylation

Synthesis of 5[β-^32^P]IP_7_, phosphorylation of proteins by CK2, and pyrophosphorylation by radiolabeled IP_7_ were conducted as described earlier (10, 24). V5-tagged mMYC or eGFP fused C-terminally to mMYC PEST domain expressed transiently in HEK293T cells was immunoprecipitated and used for 5[β-^32^P]IP_7_-mediated pyrophosphorylation as previously described (15, 23). Briefly, immunoprecipitated proteins on Protein A Sepharose beads were washed in cold PBS and resuspended in pyrophosphorylation buffer (25 mM HEPES, pH 7.4, 50 mM NaCl, 6 mM MgCl_2_, and 1 mM DTT). Pyrophosphorylation was initiated by the addition of 3–5 µCi 5[β-^32^P]IP_7_, and the reaction was incubated at 37°C for 15 min. LDS sample buffer (NP0008, Thermo Fisher Scientific) was added to the beads, and the sample was heated at 95°C for 5 min. Proteins were resolved on a 4–12% NuPAGE Bis–Tris gel (Thermo Fisher Scientific), and transferred to a PVDF membrane (GE Life Sciences). Pyrophosphorylation was detected using a phosphorimager (Typhoon FLA-9500), and protein was detected by immunoblotting. The method is explained schematically in Fig. 2A.

Back-pyrophosphorylation was conducted as described earlier (15, 23). Endogenous MYC from *Ip6k1*^+/+^ and *Ip6k1*^−/−^ MEFs was immunoprecipitated and subjected to pyrophosphorylation as described above. Radiolabeled protein as a fraction of total immunoprecipitated protein was quantified using Fiji software. The method is explained schematically in Fig. 2B.

GST-tagged hMYC and its fragments were expressed in *Escherichia coli* BL21(DE3) strain, purified on glutathione agarose beads (GE Life Sciences) using standard methods, and used for CK2-mediated phosphorylation or 5[β-^32^P]IP_7_-mediated pyrophosphorylation as described earlier (15, 24). For CK2-mediated phosphorylation, GST fusion protein on glutathione beads was incubated with 250 units CK2 (New England Biolabs) in protein kinase buffer (New England Biolabs) in the presence of 0.2 mM Mg^2+^-ATP and 1 µCi [γ^32^-P]ATP for 30 min at 30°C. For IP_7_-mediated pyrophosphorylation, GST-fusion proteins on glutathione beads were pre-treated with 250 units CK2 (New England Biolabs) in protein kinase buffer (New England Biolabs) and 0.5 mM Mg^2+^-ATP for 30 min at 30°C, then washed with cold PBS, resuspended in pyrophosphorylation buffer, and pyrophosphorylated using 5[β-^32^P]IP_7_ as described above. Proteins were detected by staining with Coomassie Brilliant Blue R250 or by transferring to a PVDF membrane (GE Life Sciences), and staining with Ponceau S dye or detection by immunoblotting. The method is explained schematically in Fig. 2D.

## Acknowledgments

The authors thank Ashok Venkitaraman for constructive and valuable feedback on this manuscript. We acknowledge R. Manorama, Manasa Chanduri, Swarna G. Thota, and Shubhra Ganguli for generating radiolabeled 5[β-^32^P]IP_7_. We thank the following for generously sharing reagents used in this work: Robert N Eisenman for HO15.19 myc^−/−^ Rat1 cell line; Maddika Subba Reddy for plasmid expressing FBW7α; Sagar Sengupta for plasmid encoding full-length human MYC. We thank members of the Laboratory of Cell Signalling for their valuable feedback. Funding: This work was supported by the Wellcome Trust/Department of Biotechnology India Alliance [WT/DBT IA, 500020/Z/09/Z], the Human Frontier Science Program (RGP0025/2016), and Centre for DNA Fingerprinting and Diagnostics core funds. A.S. is a recipient of Junior and Senior Research Fellowships from the University Grants Commission, Government of India.

## Author Contributions

P.L. and A.S. conceived, designed and performed most of the experimental work. U.C.P generated some of the expression constructs, and did some phosphorylation and pyrophosphorylation assays. V.O. performed some of the western blot experiments. P.L., A.S., and U.C.P. analyzed the data. RB conceived and coordinated the study, and analyzed the data. P.L., A.S. and R.B. wrote the manuscript with inputs from all authors. All authors agreed on the final version of the manuscript.

