## Supplemental Methods and Figures for "A pyro-phosphodegron controls MYC polyubiquitination to regulate cell survival"

**This PDF file includes:**

Supplementary Materials and Methods
Supplementary Figures S1 and S2

### Detailed methods

**Reagents**. All chemicals were purchased from Sigma-Aldrich, unless specified otherwise. The primary antibodies used in this study for immunoblotting *(*IB) or immunoprecipitation (IP), along with antibody dilution for each application, and supplier (including catalog number) are as follows: anti-MYC (Cell Signaling Technology, 5605S; IB 1:2000; IP 1:1000), anti-GAPDH (Sigma-Aldrich, G8795; IB 1:10,000), anti-α-tubulin (Sigma-Aldrich, T9026; IB 1:10,000), anti-c-Myc tag 9E10 (Sigma-Aldrich, M4439; IB 1:10,000), anti-V5 tag (Thermo Fisher Scientific, R960-25; IB 1:5000, IP 1µg), anti-GST (Invitrogen, A-5800; IB 1:10,000), anti-ubiquitinated protein FK2 (Millipore 04-263; IB 1:1000), anti-GFP (Invitrogen, A11122; IB 1:5000, IP 2µg), anti-pT58-MYC (Abcam, Ab85380; IB 1:1000). Casein kinase 2 (P6010) and restriction enzymes were purchased from New England Biolabs. Dulbecco's Modified Eagle's Medium (DMEM), Fetal Bovine Serum (FBS) and other cell culture reagents were from Thermo Fisher Scientific. NuPAGE 4-12% Bis-Tris gels were from Thermo Fisher Scientific. [$\gamma$-$^{32}$P]ATP was procured from JONAKI/BRIT.

**Expression constructs.** Full-length mouse MYC, plasmid pCX-cMyc, was a gift from Shinya Yamanaka (Center for iPS Cell Research and Application (CiRA), Institute for Integrated Cell-Material Sciences, Kyoto University, Kyoto Japan) (Addgene plasmid # 19772; GenBank Accession number NM_010849.4) (42). For expression in mammalian cells, cDNA encoding full-length mouse MYC was C-terminally tagged to the V5 epitope, and cloned into the plasmid pCDNA3.1 (+) (Invitrogen) between NheI and NotI restriction enzyme sites. For retroviral transduction, full-length mouse MYC-V5 was cloned into the EcoRI and SalI restriction sites in the plasmid pBABE-puro (gift from Hartmut Land, Jay Morgenstern, and Bob Weinberg, Imperial Cancer Research Fund, Lincoln's Inn Fields, London, United Kingdom; Addgene plasmid # 1764) (43). For expression as GST fusion proteins in *E. coli*, a plasmid encoding full-length human MYC (gift from Sagar Sengupta, National Institute of Immunology, Delhi, India) was used as a template to obtain cDNA encoding hMYC fragments corresponding to amino acid residues (aa) 201-268, and aa201-237, which were subcloned into BamHI and XhoI restriction enzyme sites, and aa238-265, which was subcloned into BamHI and NotI restriction enzyme sites in the plasmid pGEX-6P-2 (GE Life Sciences). Mutagenesis of Ser(249/250/252) to Thr, Ala, or Asp in full length mouse MYC and human MYC aa201-268 was performed by overlap extension PCR mutagenesis, and confirmed by DNA sequencing. Native and 3(S/T) versions of mouse MYC PEST domain (aa202-268) were subcloned to the C-terminus of eGFP in pT7-eGFP-C1 plasmid (Clontech Laboratories) into the BamHI and XhoI restriction enzyme sites.

**Cell culture and transfection.** *Ip6k1*$^{+/+}$ and *Ip6k1*$^{-/-}$ MEFs (22), HEK293T, and Rat1 HO15.19 *myc*$^{-/-}$ ((33)gift from Robert N Eisenman, Fred Hutchinson Cancer Research Center, University of Washington, Seattle, USA) were grown in DMEM supplemented with 10% (v/v) FBS, 1% (v/v) 1 mM L-glutamine, 100 U/mL penicillin, and 100 U/mL streptomycin. Cells were transfected using polyethylenimine (PEI) (Polysciences, 23966) using a ratio of 1:3 (DNA:PEI). All plasmids used in the transfections were purified using Plasmid Midi kit (Qiagen). Cells were harvested 48 h post-transfection for further analyses.

**Immunoprecipitation and immunoblotting.** For immunoblotting, cells were washed twice in ice-cold phosphate buffered saline (PBS), lysed in 1× Laemmli buffer, and samples were processed using standard western blot techniques. Mouse spleen samples were minced in RIPA buffer (50 mM Tris-HCl pH 7.4, 150 mM NaCl, 1% Nonidet P-40, 0.25% sodium deoxycholate, 1mM EDTA, protease inhibitor cocktail), sonicated on ice (QSONICA Q500 sonicator, 20% amplitude, 6 x 5s pulses), and lysates were centrifuged for clarification (14000 *g* for 20 min). Protein was estimated by BCA method and 100 µg total protein was taken for western blot analysis. For immunoprecipitation, cells were lysed for 1 h at 4˚C in lysis buffer (50 mM HEPES pH 7.4, 100 mM NaCl, 1 mM EDTA, 0.5% Nonidet P-40, protease inhibitor cocktail, and phosphatase inhibitor cocktail), and subjected to centrifugation at 14000 *g* for 10 min. The supernatant was precleared with 10 µL slurry of Protein A Sepharose CL-4B beads (17078001, GE Life Sciences) for 2 h at 4˚C. The specific primary antibody was added to the pre-cleared lysate, incubated overnight at 4˚C, and proteins were pulled down by incubation with 15 µL slurry of Protein A Sepharose beads for 2 h at 4˚C. Beads were washed 3 times with lysis buffer, and used for $IP_7$-mediated pyrophosphorylation or western blotting analysis. Chemiluminiscence was detected using the FlourChem E (Protein Simple) or (UVITEC Alliance Q9) documentation system. Densitometry analysis of bands was done using Fiji software.

**Half-life measurement.** Cycloheximide (CHX; 100 or 250 µg/mL, as indicated), or vehicle (DMSO, final concentration not exceeding 0.1% v/v) was added to the culture medium for the indicated time. Cells were processed as described above for immunoblotting. Protein bands were quantified, normalized to loading control and plotted against time of CHX treatment. Protein half-life was calculated from linear regression analysis of the data using GraphPad Prism 5.

**Ubiquitination assay.** Cells were treated with 20 µM MG132 for 2 h (to inhibit proteasome activity), washed twice in cold PBS, and lysed in ice-cold lysis buffer (50 mM Tris-HCl pH 7.5, 150 mM NaCl, 1% Nonidet P-40, protease inhibitor cocktail), sonicated on ice (QSONICA Q500 sonicator, 10% amplitude, 4 x 5s pulses), and centrifuged at 14000 *g* for 10 min. The supernatant was precleared by incubation with normal rabbit IgG (1 µg) for 1 h, followed by Protein A Sepharose CL-4B beads for 1 h at 4ºC. The precleared lysate was used for immunoprecipitation and immunoblotting as described above.

**Preparation of Rat1 HO15.19 mMYC-V5 cell line.** Platinum-E (Plat-E) retroviral packaging cell line (Cell Biolabs Inc.) was co-transfected with retroviral plasmid - pBABE-Puro vector, or pBABE-mMYC-V5 / mMYC-V5 3(S/T) / mMYC-V5 3(S/D) / mMYC-V5 3(S/A), and packaging plasmids, pCMV-VSV-G and pUMVC (gift from Bob Weinberg, Whitehead Institute for Biomedical Research, Cambridge, Massachusetts, USA; Addgene plasmid # 8449) (44). Media containing retrovirus particles were collected 72 h post transfection, filter sterilized through a 0.45 µm filter, and used to transduce Rat1 HO15.19 $myc^{-/-}$ cells. Polybrene (8 µg/mL) was added to the cells along with retroviral particles, and an equal amount of fresh growth medium was added after 7 h. Virus containing media were replaced with fresh growth media after 12 h, followed by a second round of infection to achieve efficient transduction. Transduced cells were selected by growth in presence of puromycin (8 µg/mL).

**Colony formation assay.** Cells were seeded in complete growth medium at low density (1500 cells/60 mm dish) in technical triplicates. After 7 days, cells were washed with PBS, fixed in presence of glacial acetic acid:methanol (1:7) for 5 min, and stained with 0.1% crystal violet in PBS for 5 min. Excess stain was removed by repeated washes with PBS, and the colonies were counted using Fiji software.

**Soft agar assay.** The soft agar assay was performed as described previously (45). In brief, cells were counted and seeded, in technical triplicates, in complete growth medium containing 0.3% top agarose over a layer of medium containing 0.5% base agarose. Fresh growth medium was applied on the surface of the top agarose every 2 days. After 3 weeks, the plates were stained with 0.01% crystal violet in 10% methanol, destained with water, and the colonies were counted using Fiji software.

**Assessment of cell viability using propidium iodide staining.** To assess cell viability upon growth in serum rich and depleted conditions, cells were cultured in either 10% or 2% FBS containing DMEM for 48 h, trypsinized, and collected in 500 µL PBS. The cell suspension was incubated with propidium iodide (2 µg/mL) for 2 min in the dark, and the percentage of dead cells, marked by uptake of propodium iodide, was assessed by flow cytometry (BD Biosciences Accuri C6 flow cytometer).

**Splenocytes preparation and proliferation assay.** Single cell suspension of splenocytes was obtained by squeezing whole spleen from either *Ip6k1*$^{+/+}$ or *Ip6k1*$^{-/-}$ mice in PBS supplemented with 2% FBS, and passing the cell suspension through a 40 µm cell strainer. Erythrocytes were depleted by hypotonic lysis in Tris-buffered ammonium chloride solution (155 mM ammonium chloride : 130 mM Tris-HCl, pH 7.6; 9:1). The remaining cells were washed twice with PBS supplemented with 2% FBS. Cells were then seeded in a 10 cm dish with RPMI 1640 medium supplemented with 10% FBS and 55 µM β-mercaptoethanol for 1 h to remove adherent cells such as macrophages and dendritic cells, and retrieve non-adherent lymphocytes. Enriched lymphocytes were seeded in a 24 well plate in RPMI 1640 medium supplemented with 10% FBS and 55 µM β-mercaptoethanol, and treated with PBS, LPS (1 µg/mL), and/or ConA (4 µg/mL) for 48 h at 37°C in a 5% humidified $CO_2$ incubator. Cell viability was assayed by incubating cells for 4 h with MTT (0.5 mg/mL). Absorbance values at 570 nm were measured using the EnSpire multimode plate reader (PerkinElmer). Absorbance values recorded in mitogen treated cells were normalized to the respective PBS treated samples. Proliferation of cells derived from *Ip6k1*$^{-/-}$ spleen was expressed as a fold change over *Ip6k1*$^{+/+}$ cells for each mitogen treatment condition.

**Statistical analysis.** Statistical analyses were performed using GraphPad Prism 5. Densitometry data for western blots were obtained using Fiji software. Band intensities of proteins of interest were normalized to the respective loading controls on the same blot. These normalized intensity values were expressed relative to the control in each blot, as detailed in the figure legends. Soft agar and clonogenic assays were conducted in technical triplicates, to obtain an average number of colonies per well for each experiment. Colonies formed by cell lines expressing mutant MYC were plotted relative to cells expressing WT MYC. *P* values are from a one-sample *t*-test, comparing mutant forms to WT mMYC expressed under the same conditions. Percentage cell death estimated by propidium iodide staining of Rat1 cell lines was compared between serum depleted and serum rich conditions using a two-tailed unpaired Student's *t*-test. Lymphocyte proliferation assay was conducted in technical triplicates for each experiment. Average absorbance value for each mitogen treatment condition was normalized to the respective mock treated sample average. Proliferation of *Ip6k1*$^{-/-}$ lymphocytes was expressed as a fold change over *Ip6k1*$^{+/+}$ lymphocytes under the same mitogen treatment condition. *P* values are from a one-sample *t*-test. $P \leq 0.05$ was considered statistically significant. The number of experimental replicates for each experiment is indicated in the respective figure legend.

### Supplemental figures

Fig. S1

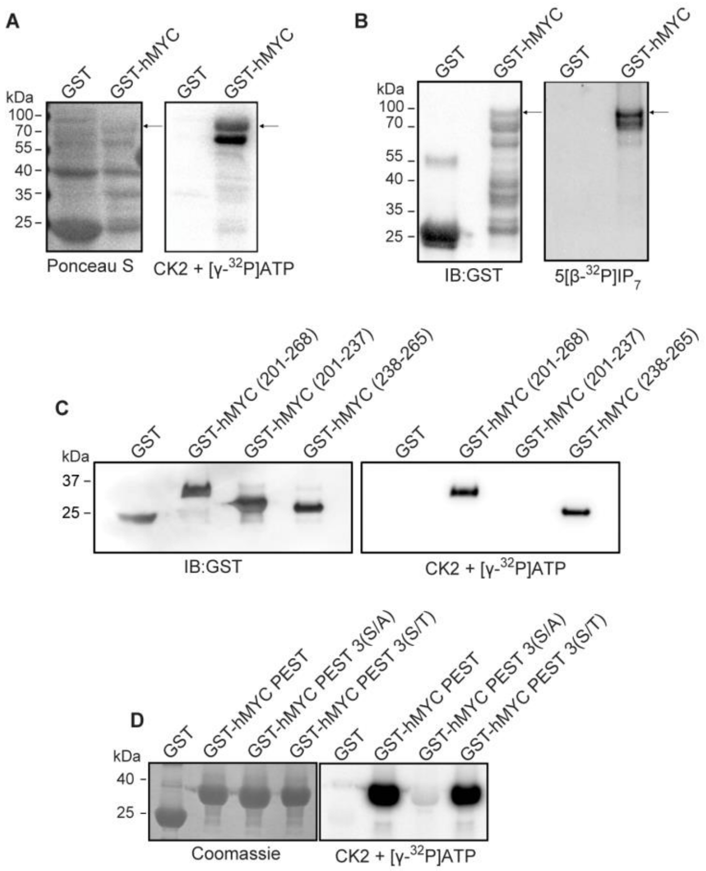


**Figure. S1. CK2 mediated phosphorylation and $IP_7$ mediated pyrophosphorylation of hMYC.** (**A**) Purified GST or N-terminally GST-tagged full length hMYC were incubated with CK2 in presence of $[\gamma\text{-}^{32}P]ATP$. Proteins were resolved by NuPAGE, transferred to a PVDF membrane, visualized by staining with Ponceau S (left), and subjected to autoradiography to detect phosphorylation (right) (N=2). (**B**) Proteins described in **A** were phosphorylated by CK2 in presence of unlabeled ATP and then pyrophosphorylated by $5[\beta\text{-}^{32}P]IP_7$ as described in **Fig. 2D.** Representative images show autoradiography to detect pyrophosphorylation (right) and immunoblotting with a GST antibody to detect protein (left) (N=2). Arrows in **A** and **B** indicate bands corresponding to GST-hMYC. (**C**) Purified, N-terminally GST-tagged hMYC fragments (start and end amino acid numbers indicated in brackets), were phosphorylated by CK2 in presence of $[\gamma\text{-}^{32}P]ATP$ as in **A**. Representative images show autoradiography to detect pyrophosphorylation (right) and immunoblotting with a GST antibody to detect protein (left) (N=2). **(D)** Purified, N-terminally GST-tagged hMYC PEST domain (aa201-268) corresponding to the native sequence, or with three Ser residues (249/250/252) mutated to Ala or Thr, were phosphorylated by CK2 in presence of $[\gamma\text{-}^{32}P]ATP$. Proteins were resolved by NuPAGE, visualized by staining with Coomassie R250 (left), and subjected to autoradiography to detect phosphorylation (right) (N=2).

**Fig. S2**

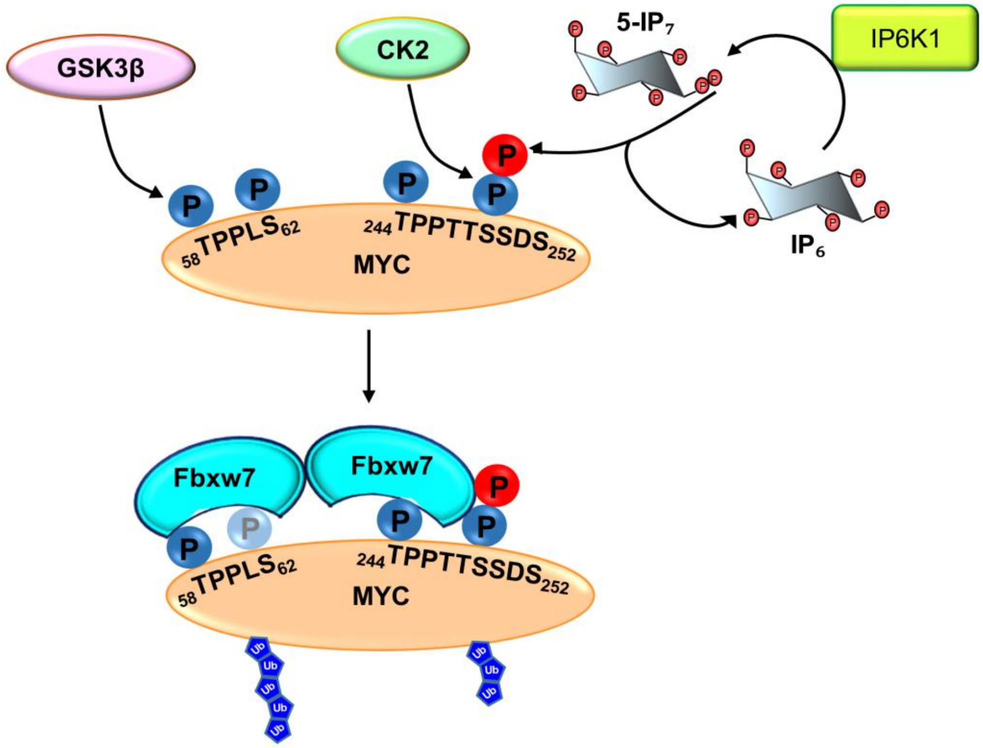


**Fig. S2. Pyrophosphorylation of MYC may enhance binding to dimeric FBW7**. FBW7 is known to interact with the N-terminus of MYC via the Thr58 phosphodegron (31). Some reports suggest that Ser62 is also phosphorylated when FBW7 binds the N-terminus of MYC (31, 39), whereas others show that Ser62 is dephosphorylated prior to FBW7 binding (18, 46). It has been shown that Thr244 is endogenously phosphorylated, and its substitution with Ala stabilizes MYC (38). Our study suggests that a phospho/pyrophosphodegron between Thr244 and Ser252 creates a second binding site for FBW7. Binding of dimeric FBW7 to two sites on MYC would stabilize their interaction, and enhance MYC polyubiquitination and degradation.
